# Single-cell Herpes Simplex Virus type-1 infection of neurons using drop-based microfluidics reveals heterogeneous replication kinetics

**DOI:** 10.1101/2023.09.18.558333

**Authors:** Jacob P. Fredrikson, Luke F. Domanico, Shawna L. Pratt, Emma K. Loveday, Matthew P. Taylor, Connie B. Chang

## Abstract

Single-cell analyses of viral infections often reveal heterogeneity that is not detected by traditional population-level studies. This study applies drop-based microfluidics to investigate the dynamics of HSV-1 infection of neurons at the single-cell level. We used micron-scale Matrigel beads, termed microgels, to culture individual murine Superior Cervical ganglia (SCG) neurons or epithelial cells. Microgel-cultured cells are subsequently enclosed in individual media-in-oil droplets with a dual fluorescent-reporter HSV-1, enabling real-time observation of viral gene expression and replication. Infection within drops revealed that the kinetics of initial viral gene expression and replication were dependent on the inoculating dose. Notably, increasing inoculating doses led to earlier onset of viral gene expression and more frequent productive viral replication. These observations provide crucial insights into the complexity of HSV-1 infection in neurons and emphasize the importance of studying single-cell outcomes of viral infection. The innovative techniques presented here for cell culture and infection in drops provide a foundation for future virology and neurobiology investigations.

## Introduction

Single-cell analyses have advanced our understanding of cellular physiology and viral infection by facilitating the observation of underrepresented phenotypes. Viral infection studies traditionally rely on population-level approaches where cells are cultured and infected in well plates (*1*). However, these approaches often overlook the heterogeneous dynamics of infection obscured by productive viral replication (*1–5*). Single-cell methods for both culture and infection provide increased insights into infectious virus production, viral replication kinetics, and genetic variability (*4–11*).

Herpes Simplex virus type 1 (HSV-1) is a ubiquitous pathogen which infects neurons to establish lifelong persistent and recurrent disease (*12, 13*). The replication, persistence, and transmission of HSV-1 are determined by the regulation and temporal expression of viral genes (*14, 15*). Single-cell studies of HSV-1 infection in epithelial cells have revealed variability in the dynamics of viral replication (*2, 16, 17*). Additionally, single-cell transcriptional analysis of HSV-1 infected epithelial cells observed a highly variable abundance of viral transcripts temporally classified as immediate-early, early, and late genes. While our understanding of epithelial cell infection at the single-cell level is improving, single-cell neuronal cell infection studies have not yet been achieved to date.

A powerful technique for studying single-cell viral infection is drop-based microfluidics (*1, 3, 18, 19*). This method generates emulsions containing monodisperse, picoliter-sized, aqueous drops suspended in oil that can be used in various single-cell assays. Drop-based microfluidic methods facilitate the encapsulation of millions of single cells, enabling in-depth, high-throughput analysis of viral and cellular heterogeneity (*1, 3, 18*). However, these methods have previously focused on infections in non-adherent cells (*18*) that were suspended in aqueous drops (*20–26*).Neurons require a soft, viscoelastic solid substrate that supports neurite development and growth, which is not compatible with the aqueous environment produced using drop-based microfluidics (*27*). Extending drop-based capabilities for adherent cultures at the single cell level would greatly facilitate understanding of viral infection in physiologically relevant cells. Support for the growth and development of cells which require solid substrates can be achieved with microscale hydrogel beads, referred to as microgels (*28, 29*). Microgels provide a homogenous and highly tunable biomimetic growth environment and have been previously used to culture multiple cell types such as neurospheres, embryonic stem cells, and induced pluripotent stem cell aggregates (*28, 29*). While microgels can offer a foundation for neuronal growth and maturation, their application in drop-based methodologies for viral infections has never been assessed. Therefore, the development of innovative techniques for individual neuron culture would facilitate single-cell studies of HSV-1 infection.

In our study, we employ drop-based microfluidics to culture and perform live-cell tracking of HSV-1 across different cell types susceptible to HSV-1 infection, including individual murine Superior Cervical Ganglia (SCG) neurons and Vero cells. The single cells are embedded in Matrigel microgels and subsequently encapsulated in drops containing defined inoculating doses of HSV-1. HSV-1 infection is visualized using a recombinant fluorescent-protein expressing reporter virus. Our results demonstrate that cells cultured within microgels are not accessible to infection, whereas cells located on the microgel surface support a greater extent of infection compared to suspension cells. Additionally, the onset of viral gene expression and replication kinetics were monitored, revealing that higher inoculating doses result in an earlier onset and progression of viral replication. In conclusion, these findings demonstrate that microgels provide a solid surface that supports neuronal growth and development, enabling productive single-cell HSV-1 infection within drops. The use of microgels for high-throughput single-cell culturing can provide a valuable tool for future research in neurobiology and virology studies, further enhancing our understanding of factors that affect viral replication dynamics.

## Results

We developed drop-based microfluidic approaches to investigate the dynamics of HSV-1 infection in individual primary neurons (Figure 1). First, individual murine embryonic SCG neurons are suspended in a Matrigel precursor solution that is processed into microgels with diameters of approximately 100 μm using a microfluidic device (Figure 1A). The microgel-cultured neurons are cultured for one week to allow for the growth and development of neurite extensions. Subsequently, neurons are infected using a co-flow inoculation device, where hydrogels and virus are simultaneously emulsified into drops. Co-flow inoculation allows precise control of the viral inoculating dose to achieve single-cell infection (Figure 1B). To visualize infection dynamics and replication kinetics, cells were infected with a dual fluorescent protein (FP) expressing reporter HSV-1. This dual-reporter HSV-1 expresses YFP driven by an immediate-early hCMV promoter and a mCherry-VP26 fusion driven by a late promoter (*30*).Detection of virus expressed YFP reports the onset of viral gene expression upon infection, while the detection of mCherry (RFP) corresponds to late viral gene expression and virion assembly (Figure 1C). To observe infection dynamics at the single-cell level, we immobilized drops on a ‘DropSOAC’ microfluidic device that enables incubation and fluorescence microscopy over the course of infection (Figure 1D) (*31, 32*). The DropSOAC allows us to monitor and analyze the progression of HSV-1 infection in individual neurons over time.

**Figure 1.**
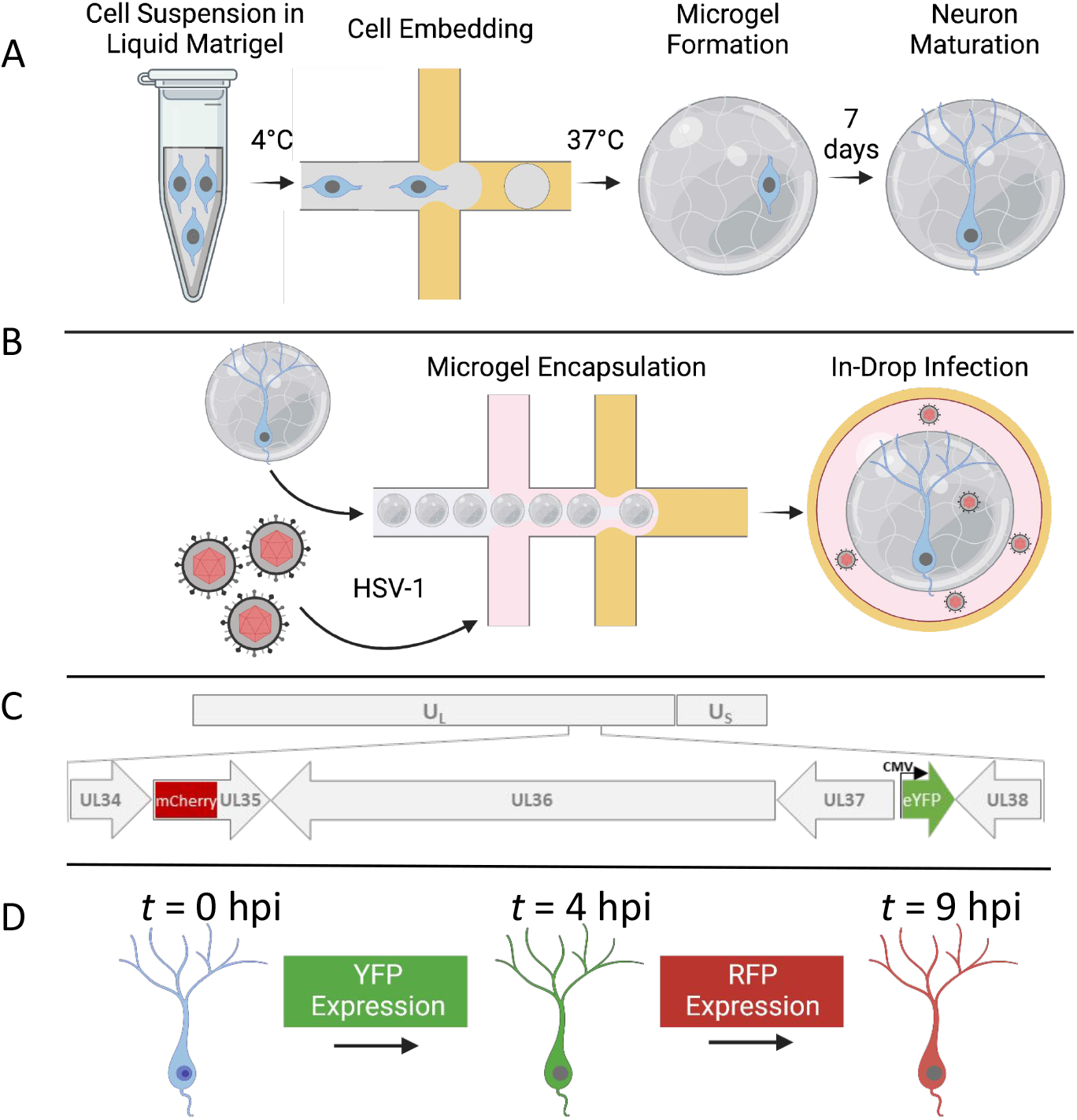
Experimental schematic for the embedding, growth, and infection of individual neurons within microfluidic drops. **(A)** SCG neurons are suspended in liquid Matrigel and emulsified in oil. The drops are incubated at 37 °C for 35 min for gelation. The microgels and neurons are washed and placed in media for 7 days for neuronal maturation. **(B)** After 7 days, the neurons are co-flowed with viral inoculum and emulsified in fluorinated oil. **(C)** A dual-fluorescent HSV-1 recombinant is employed to visualize infection. Initiation of viral gene expression is reported by YFP detection. Late gene expression is reported by RFP detection. **(D)** Individual cells can be tracked over time to observe the progression of FP detection.

### Single-Cell Infection Outcomes of Vero Cells with Dual Reporter HSV-1

To investigate the interactions between cells, microgels, and viral inoculum, we first conducted a study focusing on HSV-1 infection of Vero cells and microgels. Vero cells are an epithelial cell line commonly used for studying HSV-1 infections *in vitro* (*12*). We first evaluated whether the location of the cell in the microgel alters the likelihood of a cell becoming infected. Vero cells were either grown ‘in-microgels’, ‘on-microgels’, or placed ‘in-suspension’ before being inoculated at 10 plaque forming units (pfu) per drop using a co-flow microfluidic device (Figure 2A). YFP detection was used to evaluate the percentage of infected cells (*30*). For the in-microgel condition, Vero cells were first embedded within 100 μm microgels before emulsification with viral inoculum in drops. In-microgel infections produce only 1.7 ± 1.4% YFP positive detection at 16 hours post infection (hpi) (Figure 2B, 2C, 2F). The location of the cell in the microgel is a random event from the drop-making process, with a low percentage of cells located on the periphery of the microgel. No cells enclosed within the microgel expressed YFP (Figure 2B).Interestingly, the cells that did express YFP were all found at the edge of the microgel (Figure 2C). For the on-microgel condition, Vero cells were seeded onto prefabricated microgels and allowed to adhere for 4 hours prior to inoculation in drops. On-microgel infections produce 76.8 ± 5.0% YFP positive detection at 16 hpi (Figure 2D, 2F). For the in-suspension condition, Vero cells were emulsified in drops suspended in media with viral inoculum. In-suspension infections produce 50.0 ± 3.7% YFP positive detection at 16 hpi (Figure 2E, 2F).

**Figure 2.**
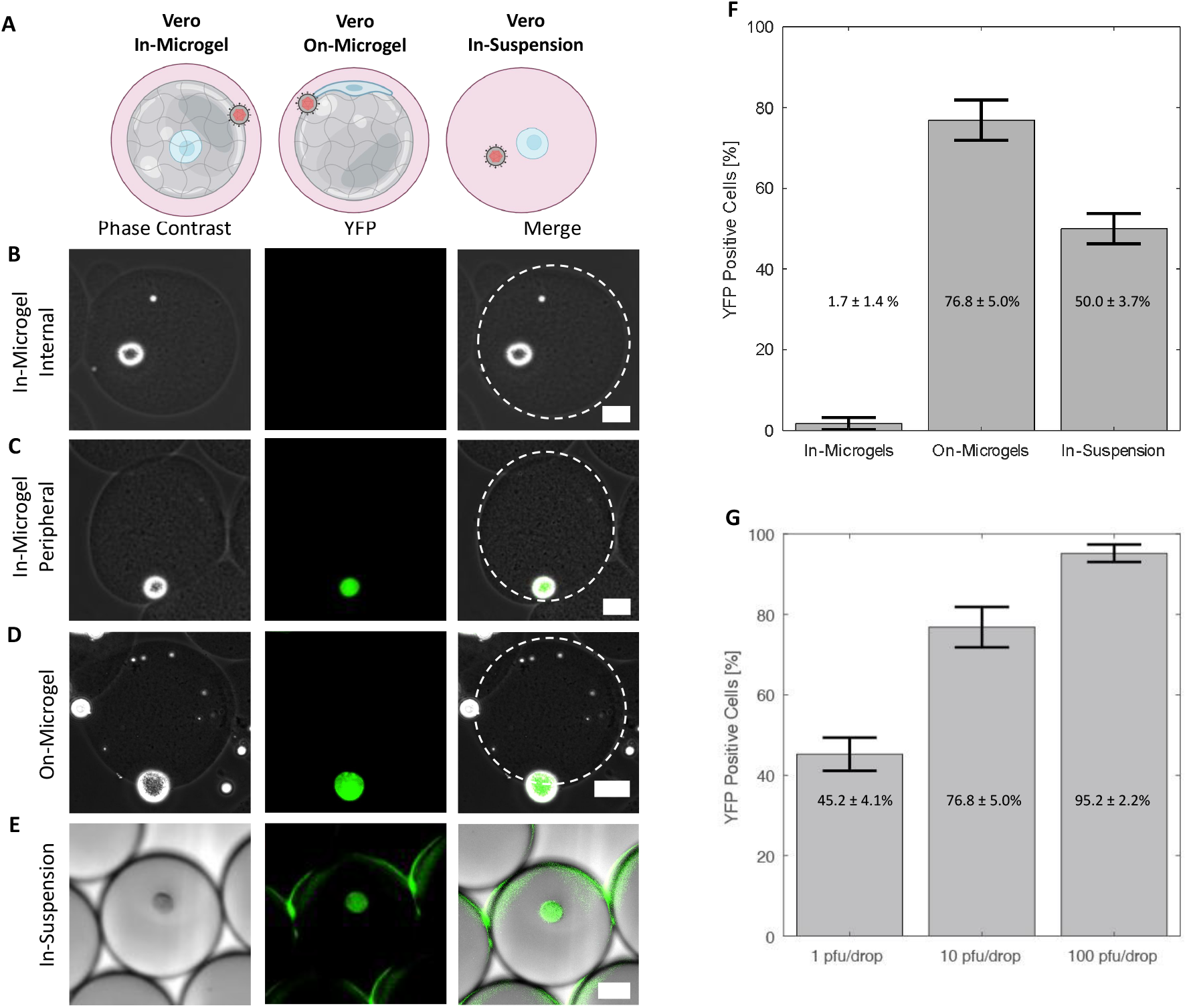
Impacts of microgels on infection. **(A)** Model of Vero cell culture in-microgels, on-microgels, or in-suspension for co-flow inoculation. **(B**,**C**,**D**,**E)** Representative phase contrast, YFP, and merged images of infected cells. Scale bars are 25 μm. **(B)** In-microgels with an internally positioned cell. **(C)** In-microgels with a peripherally positioned cell. **(D)** On-microgel. **(E)** In-suspension.**(F)** Bar graphs showing the percentage of YFP positive cells. Vero cells cultured as indicated and co-flow inoculated with 10 pfu/drop of dual reporter HSV-1. **(G)** Bar graphs showing the percentage of YFP positive cells from Vero cells cultured on microgels inoculated with 1, 10, and 100 pfu/drop. This trend was statistically significant (one-way ANOVA, *p* = 1.3 x 10-5). All infections in F and G were performed in triplicate with an average number of 200 cells per condition per replicate.

Once we determined that Vero cell cells grown on microgels were the most susceptible to infection, we further evaluated the effect of infectious dose with the co-flow inoculation system. Cells on microgels were inoculated with doses ranging from 1, 10, and 100 pfu/drop. As inoculating dose increased, the percentage of YFP positive cells increased from 45.2 ± 4.1% at 1 pfu/drop to 76.8 ± 5.0% at 10 pfu/drop and 95.2 ± 2.2% at 100 pfu/drop (Figure 2G). The extent of YFP positive cells with increasing inoculating dose was statistically significant (one-way ANOVA, *p* = 1.3 x 10-5, *df* = 8). This observation is consistent with the expectation that increasing inoculating dose leads to greater extents of infection (*33*).

We hypothesized that the low number of Vero cells embedded in microgels with detectable YFP is due to the inability of virions to penetrate Matrigel (*34, 35*). While Matrigel is a porous material, the pore size is approximately 150 nm, nearly the same size as the 150-250 nm diameter HSV-1 virion (*36*). To test our hypothesis that virion diffusion through Matrigel is limited, the interaction between mCherry-VP26 labeled virions and Matrigel was observed on a confocal microscope. Matrigel was pipetted onto glass and allowed to gel into a disc prior to the addition of mCherry-VP26 labeled virions and subsequent imaging. Over 1.5 h, we observed no HSV-1 virions diffuse past the interface of a Matrigel disc (Figure 3). Additionally, we saw no accumulation of virions at the interface, indicating that the HSV-1 particles were not adhering to or penetrating the gel and becoming immobilized (Figure 3A). These data demonstrate that HSV-1 virions likely do not diffuse through the Matrigel. To assess whether the lack of particle diffusion is related to size, we also evaluated the diffusion of fluorescent nanoparticles with average diameters of 160 nm. Like the fluorescent virions, fluorescent nanoparticles do not enter the Matrigel, but do accumulate at the aqueous interface (Figure 3B). These observations suggest that only cells which are located on the surface of Matrigel microgels are accessible to HSV-1 infection.

**Figure 3.**
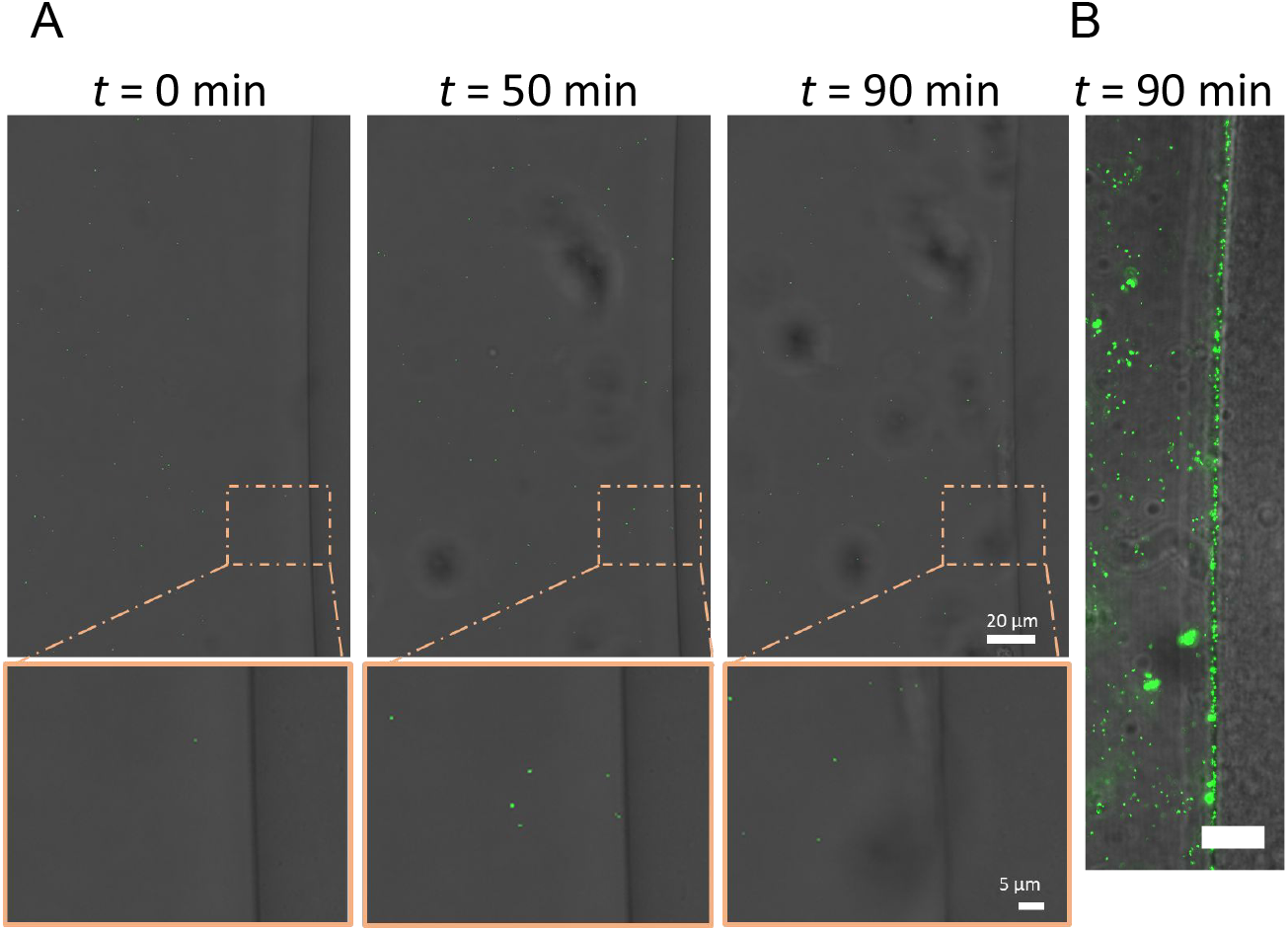
HSV-1 virions cannot enter Matrigel. **(A)** Representative images of time-lapse confocal microscopy of mRFP labeled HSV-1 virions diffusing next to a disc of Matrigel (right side of image). Images were acquired with a every 10 min for 90 min in brightfield and mRFP (false-colored green). (**B)** Nanoparticles (green) were added to the solution surrounding the Matrigel disc. Scale bar = 15 μm, *t* = 60 min.

The initial evaluations of Vero cell infections demonstrate that microgels provide a scaffold that can be used to culture and infect adherent cell lines in drop-based assays. In comparison to embedded cells, Vero cells cultured on the microgel have increased accessibility to HSV-1 infection and yield the highest percentage of detected infection. Cells that were not located on the surface of the microgels were unlikely to become infected with HSV-1 as the virions do not diffuse through Matrigel. The reduced infection observed in suspended Vero cells is hypothesized to be caused by changes to cellular permissiveness to HSV-1 infection.

### Growth of Individual SCG Neurons in Microgels

We next employed our microgel culturing system for the *in vitro* growth and development of individual primary mouse SCG neurons. Dissociated SCG neurons, unlike adherent epithelial cells such as Vero cells, require structural support to promote neurite development both during culture and microfluidic manipulation. Neurites, including axons and dendrites, are critical for neuronal homeostasis, metabolic regulation, and synaptic signaling (*27, 37*). To foster neurite development, SCG neurons were embedded in Matrigel microgels using a drop-based microfluidic device. Subsequently, the neurons were cultured to allow maturation and neurite extension over a period of seven days. After seven days in culture, the neuronal cell bodies migrated to the peripheral regions of the microgels, and robust neurites were observed either within the microgels or on their external curvature (Figure 4A). To confirm that the embedded neurons reached physiological maturity, we performed immunofluorescence staining for phosphorylated neuro-filament H (N-F), a protein localized in axons of mature neurons. SCG neurons grown in Matrigel microgels exhibited visible N-F signal in neurite extensions (Figure 4B, red). These findings indicate that microgels provide a suitable growth environment for individual neurons, enabling the production of neurite extensions and promoting maturation.

**Figure 4.**
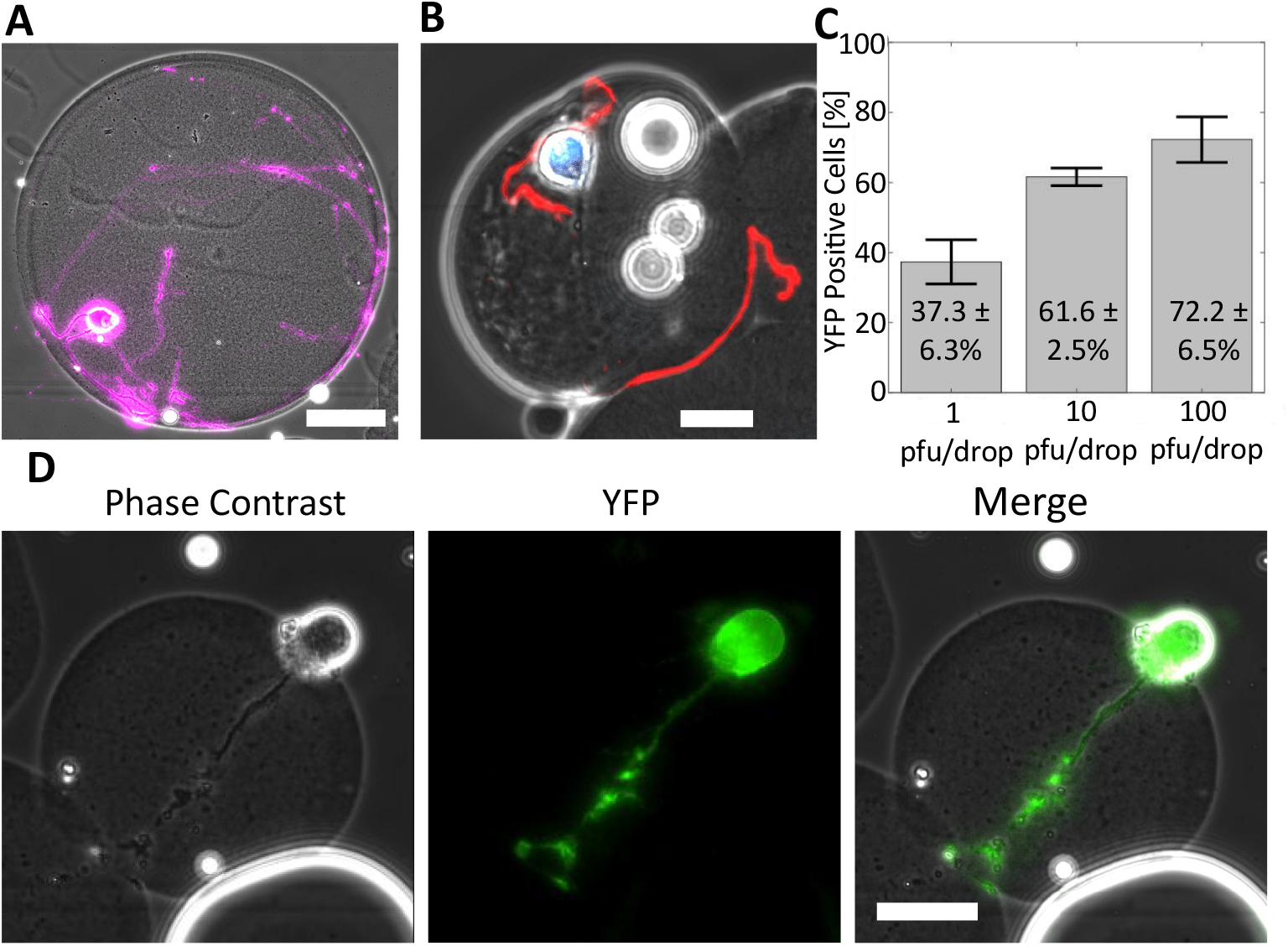
Microgels support neuronal maturation and infection. **(A)** A representative image of a mature SCG neuron in a Matrigel microgel after 7 days in culture. Cells were stained with Calcein AM, false-colored purple. Scale bar= 25 μm. **(B)** A mature SCG neuron grown in a microgel immunostained for phosphorylated neuro-filament H (Red) and Nuclei (Blue). Scale bar = 25 μm. **(C)** Bar graph showing the percentage of YFP positive neurons following infection at 1, 10, and 100 pfu/drop. Infections were performed in triplicate with an average of 158 cells per replicate per condition. Statistical significance evaluated by one-way ANOVA (*p* = 5.9 x 10^-4^). **(D)** Representative phase contrast, YFP, and merged images of an infected SCG neuron. Scale bar = 25 μm.

### Single-cell Infection of SCG Neurons

We next investigated the capacity to infect individual mature SCG neurons within microgels using dual reporter HSV-1. Microgel-cultured neurons were infected with different inoculating doses of 1, 10, or 100 pfu per drop and imaged for YFP at 16 hpi. A representative image of YFP expression in an infected neuron is shown in Figure 4D. At the lowest viral concentration of 1 pfu/drop, 37.3 ± 6.3% of SCG neurons exhibited detectable YFP (Figure 4C). SCG neurons infected with 10 and 100 pfu/drop demonstrated significantly higher percentages of infection, with 61.6 ± 2.5 and 72.2 ± 6.6% of YFP positive cells, respectively (Figure 4C). To determine the relationship between inoculating dose and YFP positivity, we conducted a one-way ANOVA and found that inoculating dose had a significant impact on YFP detection across 1, 10, and 100 pfu/drop inoculations (*p* = 5.9 x 10^-4^, *df* = 8). Our results demonstrate that primary SCG neurons cultured in microgels and infected in microfluidic drops are susceptible to HSV-1 infection and support viral gene expression, as reported by YFP detection.

### Timing and Outcomes of Viral Gene Expression in Individual SCG Neurons

We next examined the kinetics of viral gene expression in single neurons by detecting YFP for the onset of viral gene expression and RFP for late viral gene expression (*30*). To monitor and quantify the timing of viral gene expression in single neurons infected with dual-reporter HSV-1, mature microgel-cultured neurons were emulsified with viral inoculum and placed in a microfluidic chamber called a DropSOAC (Figure 5A). The DropSOAC immobilizes the drops and allows for temporal tracking and fluorescence quantification of individual cells (*31*). Images of infected neurons were acquired every 15 minutes for 16 hours, with image acquisition starting 1 hour after in-drop inoculation (Figure 5B, SI Movie 1). The onset of FP detection was determined by the point at which the fluorescent pixel intensity surpassed the background threshold value (Figure 5C).

**Figure 5.**
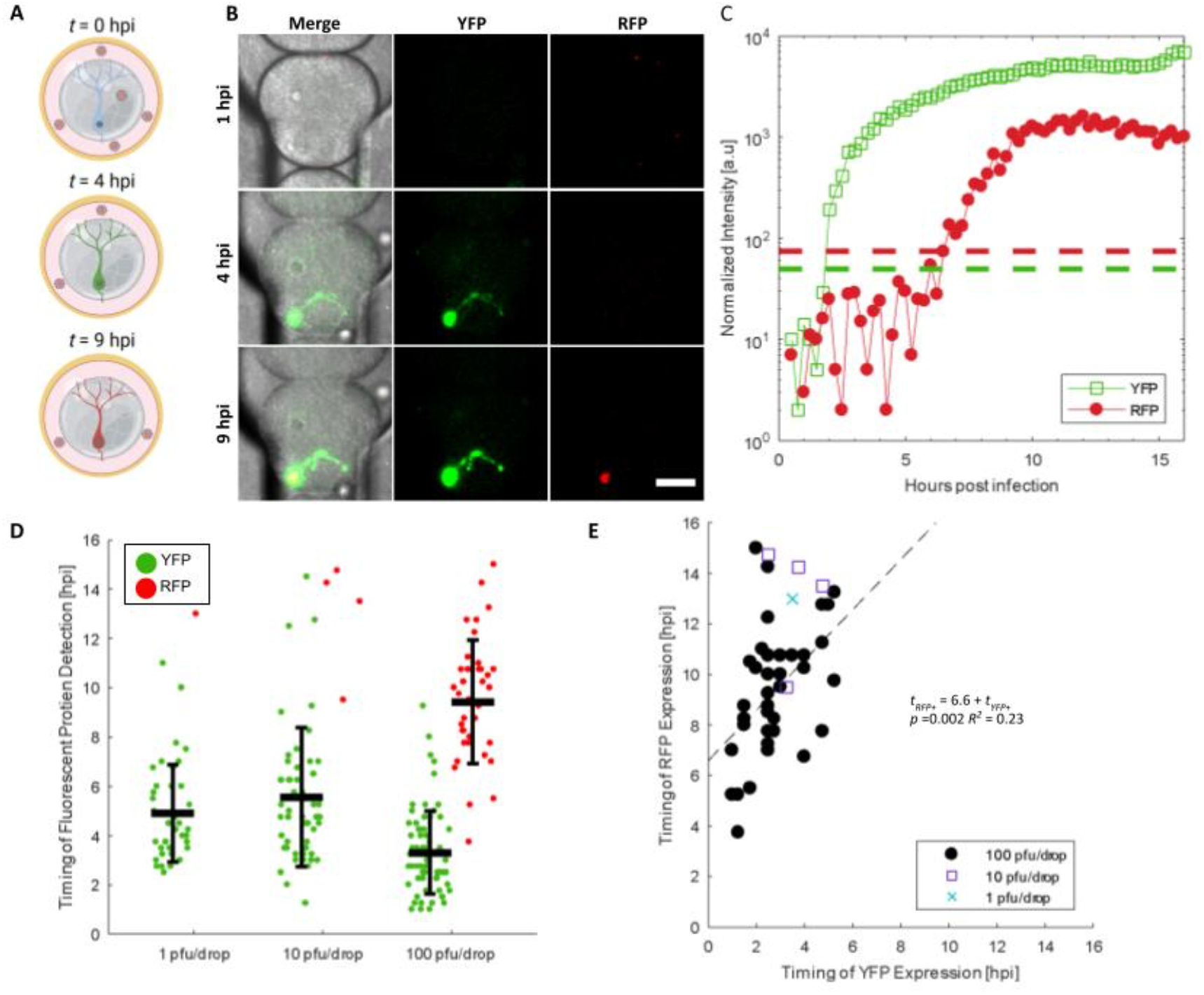
The effect of inoculating dose on neuronal infection progression and kinetics. **(A)** Schematic of experimental design for temporal tracking of YFP and RFP. **(B)** Representative images from time-lapse microscopy of an infected neuron expressing YFP and RFP in a DropSOAC chamber (*31*). Scale bar = 50 μm. **(C)** Normalized intensities of YFP (open green squares) and RFP (filled red circles) for the representative cell in (B). Dashed lines represent the threshold value above which cells are considered positive for FP detection. **(D)** Timing of YFP and RFP detection plotted with the mean and standard deviation. Each data point represents quantitation from single neurons. Statistical significance was evaluated by one-way ANOVA (*p* = 1.1 x 10^-7^). **(E)** Correlation of YFP versus RFP detection time for RFP positive neurons. (1 pfu/drop - blue X, 10 pfu/drop - purple square, 100 pfu/drop - black circle). A linear regression fit to evaluate the significance of correlation is plotted as a dashed line with the fit.

To assess the impact of inoculating dose on the detection of HSV-1 expressed FPs in single neurons, microgel-cultured neurons were infected with 1, 10, and 100 pfu/drop and imaged for 16 hours. Neurons infected with 1 pfu/drop exhibited an onset of YFP detection at 4.9 ± 2.0 hpi (Figure 5D). Neurons infected with 10 pfu/drop exhibited an onset of YFP detection at 5.5 ± 2.8 hpi. Neurons infected with 100 pfu/drop displayed the earliest onset of YFP detection at 3.3 ± 1.7 hpi. Based on a one-way ANOVA, we found that the timing of YFP detection decreased with increased inoculating dose (*p* = 1.1 x 10-7 *df* = 166), indicating that the inoculating dose significantly affects the onset of HSV-1 gene expression in single neurons.

Neurons were further examined for RFP detection, which correlates with the progression of viral replication. We observed that only 2.6 and 6.5% of YFP positive cells become RFP positive at 1 and 10 pfu/drop, respectively (Figure 5D). However, neurons infected with 100 pfu/drop exhibited much higher rates of RFP positivity, with 55.7% of YFP positive cells becoming RFP positive by 16 hpi (Figure 5D). The average timing of RFP detection in cells infected with 100 pfu/drop was 9.4 ± 2.5 hpi (Figure 5D). No cells were detected that were RFP positive and not YFP positive. These data demonstrate that in our single-neuron drop-based culturing and infection system, we can observe dose-dependent progression of HSV-1 infection in real-time.

From the time-lapse data, we estimated the progression of viral replication by calculating the timing between YFP and RFP detection in each infected neuron. At 100 pfu/drop, we observed an average time of 6.6 ± 2.2 hours between YFP and RFP detection (Figure 5E). To determine if the timing of YFP detection influences the timing of RFP detection, we implemented a linear regression model, which predicts that single neurons will become detectably RFP positive 6.6 ± 1.8 h after onset of YFP detection (*tRFP+* = 6.6(±1.8) + 1.0(±0.6)*tYFP+, p* =0.002). However, the correlation between the onset of YFP detection and conversion to YFP/RFP positive detection was weak (R2 = 0.23), indicative of the heterogenous infection in single neurons. Thus, we conclude that the timing of YFP detection is not predictive of the timing of RFP detection.

We also monitored and quantified the timing of FP detection in Vero cells infected with dual reporter HSV-1. Similar to our previous on-microgel condition, Vero cells were cultured on microgels, then encapsulated with viral inoculum, and placed in a DropSOAC device. We observed similar trends in single Vero cells to those observed in single neurons (Figure 6A). Vero cells infected with 1 pfu/drop exhibited an onset of YFP detection at 6.6 ± 2.6 hpi. Vero cells infected with 10 pfu/drop exhibited an onset of YFP detection at 5.0 ± 2.0 hpi. Vero cells infected with 100 pfu/drop displayed the earliest onset of YFP detection, at 3.5 ± 1.5 hpi. Using a one-way ANOVA, we found that the timing of YFP detection decreased with increasing inoculating doses in single Vero cells (*p* = 1.9 x 10-35, *df* = 649) (Figure 6A).

**Figure 6.**
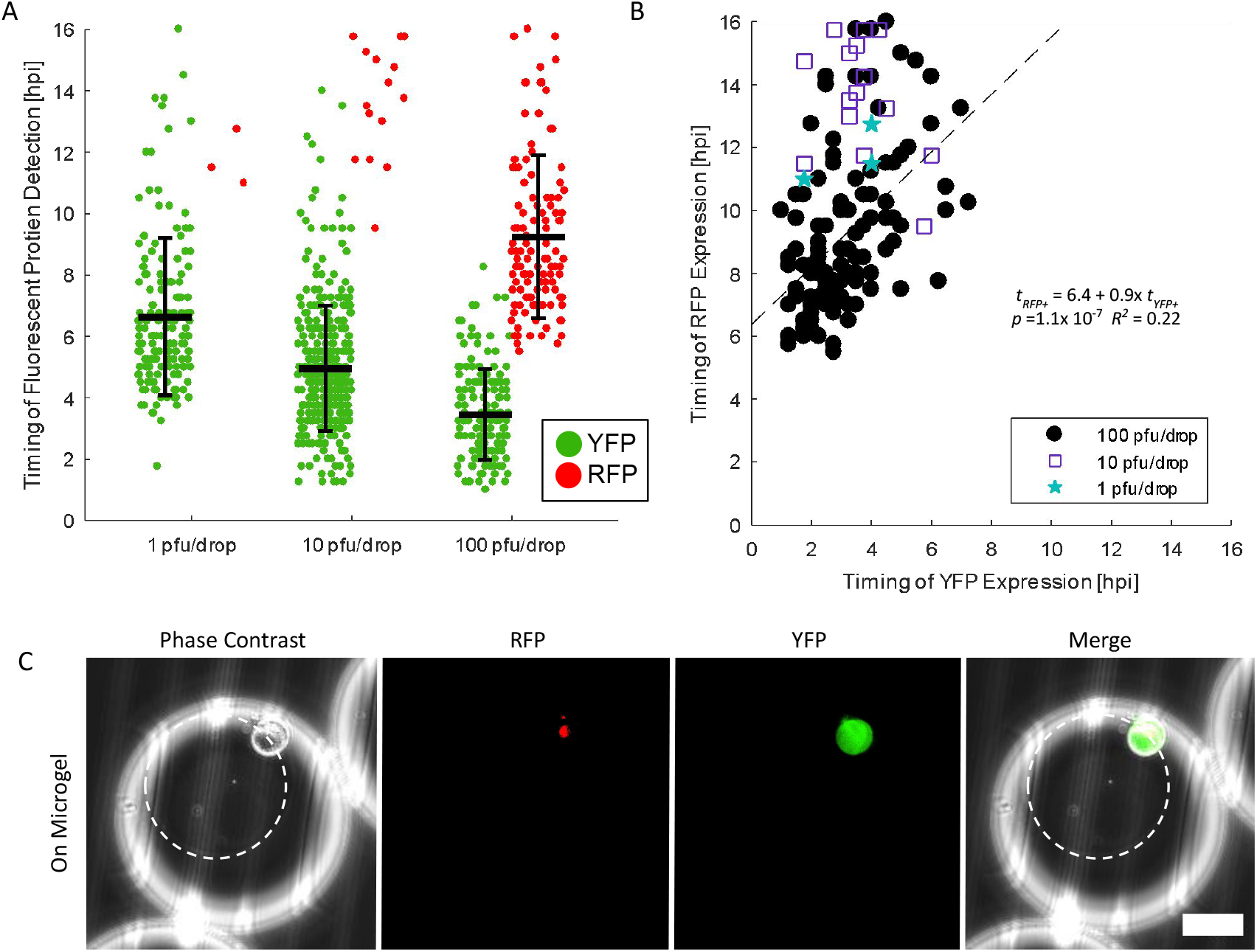
HSV-1 replication kinetics in Vero cells. **(A)** Timing of YFP and RFP expression in individual Vero cells plotted with the mean and standard deviation. Timing of YFP detection decreased with increased inoculating dose (one-way ANOVA, *p* = 1.9 x 10-35). **(B)** Correlation of YFP versus RFP detection time for RFP positive Vero cells. (1 pfu/drop - blue stars, 10 pfu/drop - purple squares, 100 pfu/drop - black circle). (Linear regression: *t*_*RFP+*_ = 6.4 + 0.9x *t*_*YFP+*_, *p* =1.1x 10-7, R2 = 0.22). **(C)** Representative images of Vero cells infected in drop on microgels. White circles outline the microgel. Scale bar = 50 μm.

In a similar trend to neuronal infection, 1.9 and 4.6% of YFP positive Vero cells became RFP positive at 1 and 10 pfu/drop, while 64.2% of YFP positive Vero cells became RFP positive at 100 pfu/drop (Figure 6A-C). From the time-lapse data, linear regression predicts that at 100 pfu/drop, Vero cells become detectably RFP positive 6.4 ± 1.0 h after onset of YFP detection; however, the fit is weak (linear regression, *tRFP+* = 6.4(±1.0) + 0.9(± 0.3) *tYFP+, p* =1.1x 10-7, R2 = 0.22) (Figure 6B). The concordance of both the extent of RFP positive cells and the progression of YFP-to-RFP detection suggests that inoculating dose, not cell type, plays an important role in determining outcomes of HSV-1 infection.

In summary, we demonstrate the use of microfluidic methods for the culture and infection of Vero cells and primary SCG neurons to observe progression of HSV-1 infection with single-cell resolution. We find that inoculating dose influences the extent and frequency of progression for HSV-1 infection in both cell types. Specifically, at higher inoculating doses, cells express detectable YFP earlier and are more likely to progress to detectable RFP progression and viral replication. These experiments validate the use of drop-based culturing and live-cell tracking of HSV-1 across different cell types susceptible to HSV-1 infection.

## Discussion

In this study, we utilized drop-based microfluidics for the culturing of individual cells and the subsequent infection with HSV-1. Culturing cells using Matrigel microgels enabled both neurons and adherent epithelial cells to maintain physiologically relevant morphologies. Following the culturing of cells within these microgels, we implemented a co-flow inoculation approach that allows for precise control over infection conditions. Subsequently, a specialized microfluidic DropSOAC device (*32*) enabled the isolation of individually infected cells and real-time observation of HSV-1 replication through detection of virally expressed FPs. Collectively, these techniques offer innovative means to culture cells and observe real-time HSV-1 replication kinetics at the single-cell level.

Neurons are the focal point of HSV-1 persistence, morbidity, and mortality; therefore, it is important to understand how HSV-1 infection progresses in primary neurons (*12*). However, primary neurons require a solid substrate that supports neurite development and growth, limiting their usability in an aqueous drop environment. To produce a solid substrate for cells, we embedded primary SCG neurons in Matrigel microgels, which promoted their maturation and facilitated microfluidic manipulation. We observed that microgel-cultured SCG neurons grew robust neurite extensions that follow the external curvature of the microgels. Additionally, immunofluorescence imaging demonstrated the presence of phosphorylated-neurofilament H in the neurite extensions, indicating SCG neuron maturation and axonal development, which was achieved by 7 days in culture. The use of Matrigel provided not only the solid substrate, but also laminin cofactors that are important for SCG survival and development (*38, 39*). This work shows that Matrigel microgels provide the necessary growth factors and support for sustained growth and viability of neurons in culture.

Microgels provide support for subsequent microfluidic manipulation of cultured neurons for in-drop infection. We achieved 72.2 ± 6.6% infection of individual neurons grown in microgels. Infections elicited a wide distribution of YFP detection onset in single neurons, ranging from 1.25 - 16 hpi. The variability could be the result of HSV-1 entry at distal axon sites leading to delays in replication. Yet, it is more likely the result of a heterogenous establishment of infection, as a wide-distribution of YFP detection onset also occurred in Vero cells. Similarly, we observed large differences in the time it takes a YFP positive cell to progress to RFP positivity in both cell types. Based on these observations, we hypothesize that the establishment and progression of HSV-1 infection is heterogeneous across individual neurons. This hypothesis is supported by recent single-cell transcriptomic studies of HSV-1 infection in epithelial cells (*2*) and neurons (*16*). Each study reported several hours of variation in the onset of viral gene expression despite synchronous inoculation. Further analyses observed that infected cells exhibit high cell-to-cell variability in late viral protein detection and abundance, suggesting that the progression of HSV-1 infection may be variable in single cells (*17*). These studies conclude that heterogeneity in HSV-1 infection is caused more by cell-to-cell variation in metabolic or immunological states within the population of susceptible and permissive cells.

Our results suggest that inoculating dose can also play a critical role in determining the productive outcome of infection. We observed that inoculating dose directly affects the kinetics of viral gene expression and the likelihood of productive replication. Our co-flow inoculation system enables precise manipulation of inoculating doses, mixed in drops with individual cells. We observed that YFP became detectable earlier in all cells infected with higher inoculating doses.

Importantly, all RFP positive neurons were detectably YFP positive before 6 hpi, suggesting that early viral transcription is more likely to elicit productive viral replication. The dose dependence on viral gene expression and productive replication aligns with other work that observed dose-dependent HSV-1 replication in human foreskin fibroblasts (*40*). Similarly, the effects of lower inoculating dose leading to slower kinetics of expression and low rates of productive replication have been observed during HSV-1 infection of other non-neuronal cells (*33, 40*). Notably, we were able to observe single-cell infection of neurons for longer observation periods. This longer observation window revealed a large population of ‘stalled’ infections. Further studies are required to understand the stage of viral replication at which progression stalls and the cellular factors that influence this outcome of infection.

To further understand the complexities of the microgel in-drop infection system, we investigated the effects of the microgel, the susceptible cell, and viral inoculum. We hypothesized that cells cultured within the microgel were inaccessible to infection, which is consistent with the minimal infection observed at 10 pfu/drop for Vero cells. Particle diffusion experiments demonstrate a lack of virion diffusion into the Matrigel. Therefore, cells cultured in-microgel are limited in their spatial accessibility to HSV-1 infection. However, cells cultured on-microgel were found to be accessible to HSV-1 and achieved a maximum of 95.2 ± 2.2% infection. The accessibility of cells to infection may partially explain why only a maximum of 78% of neurons express detectable YFP following our highest inoculating dose. The SCG neurons needed to be first encapsulated within the microgel, after which cell bodies and neurites could migrate to the gel surface. It is possible that some of the cells that did not express detectable YFP were inaccessible to infection. Alternatively, it is possible that these FP negative cells were infected but represent a population of neurons that suppressed all viral gene expression.

In conclusion, we are the first to demonstrate in-drop infection of single primary neurons cultured using microgels. The speed and scale of these microfluidic methods hold the potential for high-throughput culturing and assaying of single neurons. The microgels could also act as a scaffold for in-drop differentiation and manipulation of other primary cells (*28, 29*). While many approaches to droplet cell culture rely on suspension cells to study viral infection and replication (*3, 21, 24*), we observed an approximate 52% increase in infection when cells were cultured on microgels, compared to the same cells cultured in suspension. Finally, our drop-based single-cell approach captures heterogeneous events within a population that would otherwise be missed by bulk culturing (*3*). In conclusion, the use of microgels for high-throughput single-cell culturing can provide a valuable tool for future research in neurobiology and virology studies, further enhancing our understanding of factors that affect viral replication dynamics.

## Materials and Methods

### Vero Cell Culture

Vero cells purchased from (American Type Culture Collection, Manassas, VA) were maintained and subcultured in Dulbecco’s Modified Eagle’s Medium (DMEM) supplemented with 10% fetal bovine serum (v/v) and 1% penicillin/streptomycin in 5% CO_2_ at 37°C.

### Mouse Superior Cervical Ganglia Neuron Dissociation

Mouse Superior Cervical Ganglia (SCG) were excised from embryos at 14 days post gestation from pregnant C57Bl/6 mice. The protocol for isolating SCGs is approved by the Institutional Animal Care and Use Committee at Montana State University (protocol# 2022-52-IA). Briefly, isolated SCGs were washed with Hank’s Balanced Saline Solution (HBSS) and resuspended in 0.25 mg/mL trypsin (Gibco) in HBSS for dissociation and incubated for 15 minutes in a 37 °C water bath. Trypsinized SCGs were centrifuged and resuspended in 1 mg/mL trypsin inhibitor (Gibco) in HBSS, then incubated for 5 minutes in a 37 °C water bath. Next, SCGs were centrifuged and resuspended in complete neurobasal (neurobasal media (Gibco), 1X B27 (Gibco), 60 ng/mL 2.5S NGF (Millipore Sigma) and 1% penicillin/streptomycin + glutamate (Gibco)) and dissociated by trituration using a 5 mL Pasteur pipette (*41*). Dissociated neurons were then cultured as described in complete neurobasal media.

### Dual Reporter Herpes Simplex Virus-type 1

Dual reporter HSV-1 was constructed, isolated, and characterized as previously described (*30*). Vero cells were used for viral stock production and plaque assay estimation of viral titers.

### Microfluidic Device Fabrication

Negative master molds for the microfluidic devices were prepared using standard photolithography techniques (*42*). Negative master molds were made with Nano SU-8-100 photoresist (Microchem, Round Rock, TX, USA) on 3 silicon wafers (University Wafer Inc., Boston, MA, USA University Wafer ID: 447). The microgel drop-maker and the suspension cell co-flow inoculating device were fabricated to be 100 μm tall. The co-flow microgel inoculating drop-maker and the DropSOAC (*31*) chambers were fabricated to be 150 μm tall. Devices were treated with (tridecafluoro-1,1,2,2-tetrahydrooctyl) trichlorosilane (1% v/v) (Gelest) in fluorinated oil HFE 7500 (3M, Saint Paul, MN, USA) and left for solvent evaporation at 55 °C.

### Microgel Production and Cell Encapsulation/Seeding

Matrigel microgels were produced through drop-based microfluidics using previously established protocols (*43*). Briefly, liquid Matrigel at 4°C and 1.5% w/w fluorosurfactant (008, RAN Biotechnologies, MA, US) in HFE 7500 were loaded into luer lock syringes and injected into the 100 μm drop-maker using syringe pumps. The flow rates used were *Q*_*Matrige*l_ = 100 μL/h and *Q*_*HFE*_ = 900 μL/h. All equipment and reagents were refrigerated at 4 °C to prevent premature gelation of the Matrigel. Drops were collected in microcentrifuge tubes and incubated at 37 °C for 35 min to gel the drops. The resulting microgels were washed with equal volumes of 1H,1H,2H,2H-Perfluoro-octanal (PFO) - HFE 7500 (20% v/v) and PBS to drops. The cleaned microgels were then collected in PBS.

In experiments where Vero cells or dissociated SCG neurons were encapsulated in microgels, cells were suspended in the liquid Matrigel at 1 x10^6^ cells/mL prior to drop making. After collecting the resulting microgels in PBS, they were placed in well plates with the appropriate growth medium. Vero cells were maintained in DMEM - 10% FBS - 1% penicillin/streptomycin in 5% CO_2_ at 37 °C for 4 h before experimentation. SCG neurons were grown in complete neurobasal media at 37 °C in a 5% CO_2_ enriched atmosphere for 7 days prior to experimentation to allow maturation and neurite growth. 24 h after encapsulation, SCGs were supplemented with 1 μM Cytosine-β-D arabinofuranoside (Sigma C6645), a compound cytotoxic to mitotically active, non-neuronal cells. In experiments where Vero cells were seeded onto the microgels, empty microgels were mixed with 1x10^6^ cells per 1 mL_Microgel_ in a well plate. Cells were allowed to adhere for 4 h prior to experimentation.

### Immunofluorescence Staining and Imaging

After 7 days in culture, neurons grown in microgels were collected in microcentrifuge tubes and washed with PBS. Cells were fixed with 1% glutaraldehyde for 1 min and washed 3x with PBS. Cells were blocked with 2% BSA/PBS at room temperature for 1 h and washed 3x with PBS. Cells were permeabilized with 0.1% Triton-X / 0.125% BSA/PBS and washed 3x with 0.125% BSA/PBS. The primary antibody, anti-Phosphorylated Neurofilament H (NF-H) (BioLegend), was added at 5 μg/mL, incubated at room temperature overnight, and then washed 3x in 0.125% BSA/PBS. The secondary antibody, Goat anti-Mouse IgG (H+L), DyLight 550 (#84540 ThermoFisher), and Hoechst 33342 Solution (ThermoFisher) were added at 5 μg/mL and 20 μg/mL respectively, for 1 h at 37 °C and then washed 3 times with 0.125% BSA/PBS. Cells and microgels were loaded onto glass slides and imaged on an epi-fluorescent microscope (Nikon Ti2). Cells were imaged in Phase Contrast/DAPI/RFP.

### In-Drop Infection Procedures

Cells were infected in drops. To infect cells seeded onto or cultured in microgels, microgels were collected from well plates and pelleted by centrifuging the microgels at 200 x *g* for 1 min. The pelleted microgels were loaded into luer lock syringes. HSV-1 inoculum was diluted into the appropriate media. For Vero infections, HSV-1 stock was diluted in DMEM - 10% FBS - 1% penicillin/streptomycin. For SCG infections, HSV-1 stock was diluted in complete neurobasal media. HSV-1 was diluted at concentrations of 1.1 x 10^6^, 1.1 x 10^7^, and 1.1 x 10^8^ pfu/mL, resulting in inoculating conditions of 1, 10, and 100 pfu/drop, respectively. The virus solutions and a 1.5% solution of RAN in HFE were loaded into individual luer lock syringes. The three syringes were loaded onto syringe pumps and injected into the appropriate inlet channels of the microfluidic co-flow microgel inoculation device. Flow rates were *Q*_*HFE*_ = 2500 μL/h, and *Q*_*Matrigel*_= *Q*_*Virus*_= 250 μL/h. Drops were collected into microcentrifuge tubes and either placed in an incubator at 37 °C for end-point imaging or injected into DropSOAC chambers for time-lapse imaging.

To infect Vero cells suspended in media, Vero cells were removed from subculture using Trypsin-EDTA and washed in PBS. Cells were suspended in DMEM - 10% FBS - 1% penicillin/streptomycin at 1x10^6^ cells/mL and loaded into a luer lock syringe. HSV-1 was diluted into DMEM - 10% FBS - 1% penicillin/streptomycin at a concentration of 3.8 x 10^7^ pfu/mL resulting in an inoculating condition of 10 pfu/drop. The virus solution and a 1.5% solution of RAN in HFE were loaded into individual luer lock syringes. The three syringes were loaded onto syringe pumps and injected into the appropriate inlet channels of the co-flow suspension cell inoculating drop-maker. Flow rates were *Q*_*HFE*_ = 2000 μL/h, and *Q*_*Cells*_*= Q*_*Virus*_ = 250 μL/h. Drops were collected into microcentrifuge tubes and placed in an incubator at 37 °C for endpoint imaging.

### End-Point Imaging of Inoculated Cells

To visualize in-drop infection, drops were loaded into capillary tubes and imaged. Cells were imaged in phase contrast/FITC/TRITC. To quantify the percentage of infected cells at 16 hpi, the drops containing infected cells were broken using 20% v/v PFO-HFE. Breaking the emulsion allowed for easier visualization and quantification of cells. The broken supernatant containing infected cells was pipetted onto a polytetrafluoroethylene (PTFE) printed microscope slide and imaged.

### Time-Lapse Imaging of Inoculated Cells

To track the progression of FP detection in single cells, drops containing infected cells were loaded into DropSOAC devices with modified aluminum capsules (*31*). The capsules were placed in a microscope stage top incubation chamber (OKOlab) at 37°C. Images in Phase Contrast/YFP/RFP were taken every 15 min for 16 h. Tile scans of each chamber were taken to capture as many cells as possible. Image acquisition began within 1 hr post-inoculation.

### HSV-1 Diffusion with Matrigel Experiments

To evaluate HSV-1 virion diffusion through Matrigel, a time-lapse imaging series of virions interacting with the Matrigel interface was performed via inverted laser scanning confocal microscopy (iCLSM) (Stellaris DMI8, Leica). Matrigel (20 μL) was pipetted onto a 35 mm glass-bottom dish (MatTek) and gelled at 37 °C, forming a hemisphere of solid gel on the glass surface. PBS was added to the glass-bottom dish to submerge the Matrigel. The PBS and Matrigel-containing dish was placed on the microscope stage, and a time-lapse imaging series (XYT) was initiated, with the focal point centered on the Matrigel-PBS interface. At *t* = 0 min, 1 x 10^7^ pfu of mRFP-VP26 tagged virions or 1 x 10^8^ of Yellow-Green fluorescent tagged nanoparticles (160 nm, FluoSpheres™ Carboxylate-Modified Microspheres Catalog #F8811, Thermo Fisher Scientific) were added to the dish. Images were acquired at 63x every 10 min for 90 min in two channels: a transmitted light channel (brightfield) and a fluorescence channel (mRFP, FITC).

## Statistical Analysis

### Percentage of YFP Positive Cells

To quantify the percentage of YFP positive cells, end-point images were analyzed using Fiji ImageJ (*44*). Cells were found in brightfield. The background intensity of the YFP channel was subtracted from the image, and the percentage of YFP positive cells was quantified. Experiments were repeated in triplicate. Cell counts for Vero cells embedded in microgels ranged from 79 -280 cells/replicate, for Vero cells grown on microgels from 165 - 390 cells/replicate, for Vero cells in suspension from 228 - 456 cells/replicate, and for SCG neurons from 84 - 209 cells/replicate. Multiple comparison tests and one-way ANOVAs were performed in Matlab.

### The Timing of YFP and RFP Detection

To quantify the timing of YFP and RFP detection, time-lapse images were analyzed using Fiji ImageJ (*44*). Cells were difficult to identify in brightfield, so cells were identified by finding YFP positive cells at 16 hpi. Once located, a 30 μm diameter circular region of interest (ROI) was drawn around each individual cell. The maximum pixel intensity in YFP and RFP for the ROI was then measured for each time-frame. The frame 1 maximum pixel intensity of each ROI was subtracted from the subsequent frames for that ROI. The noise threshold was found to be 50 arbitrary units (a.u.) for YFP and 75 a.u. for RFP. The timing of YFP and RFP detection were defined as the first time-frame the YFP and RFP pixel intensities in each ROI were greater than the noise threshold for two consecutive frames. For SCG time-lapse studies, experiments were repeated 3-4 times with a total of 174 cells analyzed. For Vero timelapse studies, experiments were repeated 4 times with a total of 669 cells analyzed. One-way ANOVAs and linear regressions were performed in Matlab.

## Funding

This work was supported by:

National Institutes of Health grant 1R56AI156137-01 (C.B.C.)

National Institutes of Health grant R21AI146952-01 (M.P.T.)

National Institutes of Health grant R21AI171724-01A1 (M.P.T.)

Montana Agricultural Experiment Station HATCH project MONB00029-7004782

(M.P.T.)

National Science Foundation CAREER grant 1753352 (C.B.C).

## Author contributions

Conceptualization: JPF, LFD, EKL, MPT, and CBC Investigation: JPF, LFD, and SLP

Visualization: JPF and LFD Supervision: MPT and CBC Writing—original draft: JPF and LFD

Writing—review & editing: JPF, LFD, EKL, MPT, and CBC

## Data and materials availability

All data are available in the main text or the supplementary materials. The original imaging files, other data, analysis files, device designs, and reagents are available, upon request, and may be subject to materials transfer agreements (MTAs).

## Supplementary Materials

Supplemental Movie 1

